# Dorsolateral prefrontal cortex drives strategic aborting by optimizing long-run policy extraction

**DOI:** 10.1101/2024.11.28.625897

**Authors:** Jean-Paul Noel, Ruiyi Zhang, Xaq Pitkow, Dora E. Angelaki

## Abstract

Real world choices often involve balancing decisions that are optimized for the short-vs. long-term. Here, we reason that apparently sub-optimal single trial decisions in macaques may in fact reflect long-term, strategic planning. We demonstrate that macaques freely navigating in VR for sequentially presented targets will strategically abort offers, forgoing more immediate rewards on individual trials to maximize session-long returns. This behavior is highly specific to the individual, demonstrating that macaques reason about their own long-run performance. Reinforcement-learning (RL) models suggest this behavior is algorithmically supported by modular actor-critic networks with a policy module not only optimizing long-term value functions, but also informed of specific state-action values allowing for rapid policy optimization. The behavior of artificial networks suggests that changes in policy for a matched offer ought to be evident as soon as offers are made, even if the aborting behavior occurs much later. We confirm this prediction by demonstrating that single units and population dynamics in macaque dorsolateral prefrontal cortex (dlPFC), but not parietal area 7a or dorsomedial superior temporal area (MSTd), reflect the upcoming reward-maximizing aborting behavior upon offer presentation. These results cast dlPFC as a specialized policy module, and stand in contrast to recent work demonstrating the distributed and recurrent nature of belief-networks.

## Introduction

In dynamic environments animals continuously select actions to maximize a utility function^1^. Namely, we select actions in an attempt to balance the acquisition of reward, now or in the foreseeable future, with the expenditure of resources^2^ (e.g., energy). To enable adaptive behavior, these policies driving action selection ought to be sensitive to changes in the value of available options, our uncertainty about the state of the world, and/or our ability to alter the environment. How the brain instantiates policies—e.g., deciding what actions are worth an effort —is not fully understood. In particular, we do not fully understand how strategic, long-term planning (e.g., beyond a single trial, or offer) is updated and maintained within dynamic, closed action-perception loops.

Significant progress toward answering this question has come from reinforcement learning^3^ (RL) and the development of artificial neural network agents. In this setting, we can construct various network architectures, define utility functions, and examine learned policies. This approach, when applied to neuroscience, first led to the conjecture that dopamine-driven plasticity in the striatum translates experienced action-reward associations into optimized behavioral policies^4^. More recently, it has been suggested that a second neural RL algorithm exists, a meta-reinforcement learning system that is bootstrapped yet independent from the dopaminergic striatum^5, 6^. This latter meta-RL network is hypothesized to (1) be reflected in neural activity (as opposed to synaptic weights) of the pre-frontal cortex, (2) be more model-based than the dopaminergic striatum^7^ (but see^8^), and (3) more readily allow for generalization^9^ (see^5^ for further detail).

Prior work has largely supported claims casting the prefrontal cortex as implementing components of an RL algorithm. For instance, this area (or sub-divisions within) reflect(s) subjective value^10^, decision confidence^11, 12^, the effort required to harvest a reward^13, 14^, and/or task strategy selection^15, 16^. These seminal studies, however, have important limitations: (1) most (primate) studies artificially dissociated periods of action, perception, and reward, (2) they typically only allow for small and discrete state-spaces (e.g., two-alternative forced choice), and (3) often involve trading an immediate uncertain high-value reward for an assured, but also immediate, low-value outcome. In other words, many tasks classically used to probe macaque prefrontal cortex neurophysiology do not unequivocally need model-based RL. In fact, these tasks are not always modeled as an RL problem, and there is no evidence that they require among other, generalization^5^, long-term planning^17, 18^, and the forgoing of immediate reward for a better long-term return.

In an effort to study real-world computations, our group has developed a closed-loop task wherein macaques continuously navigate a virtual environment and stop at the location of transiently presented (300ms) targets— akin to catching a briefly flashing firefly^19, 20^ (**Fig. 1A**). This task is executed in continuous state and action spaces, requires an evolving belief (i.e., posterior probabilities) over partially observable states, and is learned via sparse rewards (i.e., trials may last many seconds and animals are only rewarded at the end of an action sequence). Importantly, within this naturalistic context, animals generalize with no further training to variants of the task requiring novel sensorimotor mappings, inference, decision-making, and foraging^21^. That is, within this task, animals demonstrate the use of a model-based approach and zero- or one-shot generalization^21^. Further, we have previously developed actor-critic RL networks instantiating artificial agents performing this “firefly task.” We found that modular architectures — separating the computation of belief from action (in the actor), and/or belief from value (in the critic) — help agents generalize quickly within this task^22^ (also see^23^ for a similar finding).

**Figure 1.**
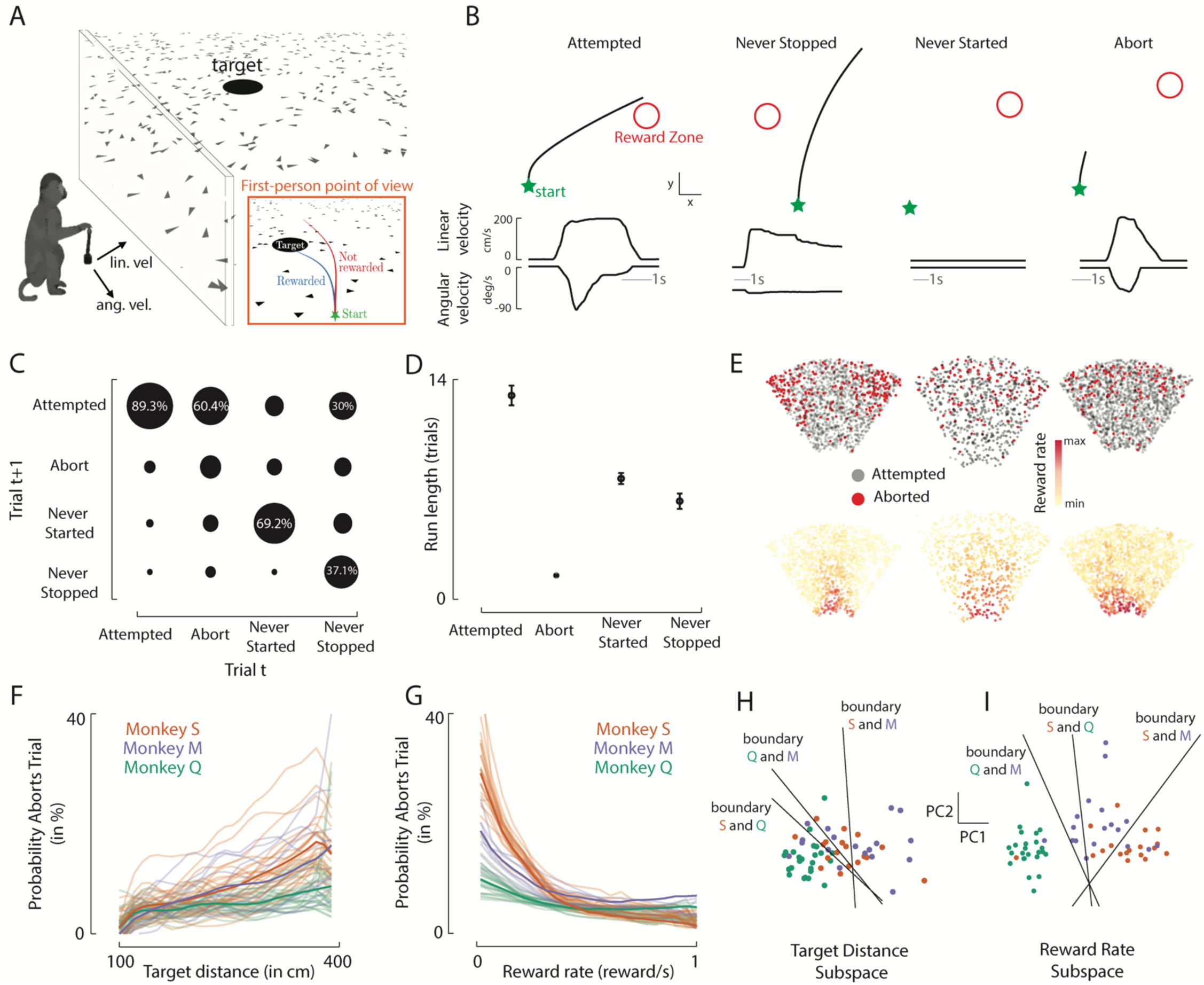
Macaques abort trials to maximize session-long reward rates. **A.** Schematic of the task and reward structure. Animals use a joystick controlling their linear and angular velocity to path integrate to the location of a briefly presented (300ms) target. Targets are presented within 100 to 400 cm of depth (radial distance), and −40° to +40° eccentricity (uniform distribution). Animals are rewarded for stopping within 65 cm of the target. Panel shows both an allocentric point of view (3d) and as an inset (orange) a rendering of the first-person point of view of the macaque **B.** Example trials showing stereotypical “failure modes”: attempted yet unrewarded trials (leftmost), trials where the animal did not stop within the maximum trial time (7 seconds, “never stopped”, 2^nd^ column), trials where the animal never started moving (3^rd^ column), and trials where the animal accelerated and then abruptly aborted the trial (rightmost). Bottom of each column shows a time-series of linear (in cm/s) and angular (deg/s) velocity (gray inset shows 1 second for each example trial). Top of each columns show a bird’s-eye-view. **C.** Transition probability matrix between trial types (trial *tr* along the x-axis and *tr* + 1 along the y-axis). Circle size is proportional to probability of transition. **D.** Run lengths of a particular trial type. Error bars are ± 1 standard error of the mean (s.e.m.). **E.** Three example sessions. Top: Location of targets (overhead view) that were attempted (gray) or aborted (red). Bottom: same plot as above, while plotting the reward rate offered by a given targets (according to animal- and session-specific likelihood of obtaining reward for that target, and the time required to reach it). **F.** Probability of aborting trial as a function of target distance (integrated sums to ‘1’). Each animal is plotted in a different color; opaque line shows the average for the specific animal, while translucent lines show individual sessions. **G.** Same as **F,** plotted as a function of reward rate. Animals are less likely to abort trials that offer a high reward rate. **H.** Data from **F** is plotted in its two leading principal components (colored as a function of animal) and linear discriminant analysis is applied to classify sessions by animal (black lines show bifurcation lines). **I.** Same as **H** when projecting data from **G.**

Here, we build on this approach both experimentally and computationally to examine (1) if and how macaques employ long-term strategic plans within this closed-loop task, (2) what algorithms or RL architectures they may use for implementing long-term policies, and (3) what neural codes may subserve these session-long policies.

## Results

### Macaques reason about their own performance in maximizing reward rate

In prior work we have described macaques’ mean performance^20, 21^ and their neural correlates^24–26^ in successfully navigating toward and stopping at the location of briefly presented targets (“fireflies”) in virtual reality. The animals are rewarded on 61.5% of trials. Here, instead, we examine an equally important (and still numerous, 38.5%) set of trials: “failure modes” within this task. These are the trials that are arguably the most informative regarding the animals’ strategies. We particularly focus on examining long-term policies (which may be sub-optimal on a shorter time-frame), and changes in this policy given matched experimental offers.

Examining the radial distance animals travelled as a function of the presented target distance (uniform distribution, 1 to 4 meters in radial distance, see *Methods*) revealed four categories of unrewarded trials (see distribution for an example session in **Fig. S1**). The majority of these were attempted trials: the animals simply misestimated—typically undershot^20, 21, 24, 25, 26^—the location of targets (52.6% of unrewarded trials; **Fig. 1B** leftmost). In another two categories of unrewarded trials, the animals either never started navigating toward the target (13.5% of unrewarded trials; **Fig. 1B** second column), or never stopped (trial timed-out; 17.4% of unrewarded trials; **Fig. 1B** third column). Lastly, on 16.5% of unrewarded trials (average of 114.3 trials per session), the animals started navigating toward the target (linear velocity threshold > 10 cm/s; minimum distance travelled > 5 cm) and then abruptly stopped, thus aborting the trial (maximum distance travelled < 30% target distance; **Fig. 1B** rightmost).

The aborted trials appeared to be a deliberate task strategy, in contrast to the “never started” or “never ended” trials which are seemingly better described as disengaged states^27, 28^. Namely, the transition probability between trial-types (**Fig. 1C**) showed that after an attempted trial (rewarded or not), animals most often attempted the next trial (89.3% of trials). Similarly, animals disproportionately repeated “never started” trials (69.2%) or “never stopped” trials (37.1%). In contrast, after an *aborted* trial, the animals most often attempted the next trial (60.4%) rather than aborting again (22.1%, Chi-square test, χ^2^ = 19.9, *p* = 2.9 × 10^"1$^). This same pattern can be observed by computing the number of consecutive trials of the same category (run-lengths, **Fig. 1D**). Animals on average attempted 13.9 trials in a row (± s.e.m=0.6). Similarly, “never started” and “never ended” trials had relatively long run lengths, respectively of 7.6 (± 0.34) and 6.0 (± 0.47) trials. In contrast, the average run length of aborted trials was only 1.2 trials (± 0.05), suggesting that this behavior was intentional and not indicative of a longer-term disengagement state^27, 28^. Analysis of eye movements also supported this conclusion: on aborted trials animals detected and saccade toward the target just as they did in attempted trials (**Fig. S2**), yet they chose to abort the trial.

The experimentally-imposed variable that most clearly drove aborting behavior was the radial and angular distance of the presented target (**Fig. 1E**; three example sessions shown). However, there was a large spatial overlap between trials animals chose to attempt (gray), and those they chose not to (red, **Fig. 1E**, see “salt-and-pepper” mixing of attempted and aborted trials). In other words, the animal’s policy (e.g., attempt or not) could differ for a “matched offer” (i.e., same location of target). In turn, we hypothesized that an important factor determining whether animals aborted or not trials was the afforded reward rate of a given trial. The afforded reward rate, however, is not an experimentally-imposed variable: it depends on the animal- and session-specific probability of being rewarded for a given target presentation, and on how fast the animal completes the trial. Thus, if aborting behavior is driven by afforded reward rates, this behavior is animal- and session-specific.

We computed the reward rate afforded by targets in an animal- and session-specific manner (see *Methods*), and expressed the probability of aborting a trial as a function of either target distance (experimentally-imposed, **Fig. 1F**) or afforded reward rate (a variable determined in part by the macaque itself, **Fig. 1G**). Monkeys aborted more trials as the target distance (at presentation) grew, but the exact nature of this relationship varied across sessions (**Fig. 1F**), mimicking the variance observed on individual trials (**Fig. 1E**). Instead, when aborting probability is expressed as a function of reward rate (**Fig. 1G**), each session could be nicely attributed to a particular animal and it becomes apparent that monkeys were preferentially aborting trials that offered a low reward for the time investment required, To quantitatively capture this pattern, we projected the probability of aborting trials as a function of distance (**Fig. 1F**) and reward rate (**Fig. 1G**) into their first two principal components (respectively accounting for 89.9% and 91.3% of the total variance). In these spaces, we performed linear discriminant analysis (LDA^29^) to classify held-out (10-fold cross-validation) sessions as belonging to one of three monkeys (chance ∼ 36.6% given the unequal number of sessions per animal). The classifier based on target distance achieved an accuracy of 70.0% (± 0.8%; **Fig. 1H**), while the classifier based on reward rates achieved an accuracy of 89.6% (± 0.7%; *p* < 0.001; **Fig. 1I**; 28% better than the classifier based on target distance).

Together, these results show that macaques employed the strategy of deliberately aborting trials (forgoing reward on trial *tr*) to increase their reward rate over the long run. This behavior was animal- and session-specific, demonstrating their ability to take into account their own performance (in terms of probability correct and trial duration) and implement an adaptive long-term (∼ session-long) plan.

### Modular RL actors with access to specific values from critic strategically abort trials to maximize reward rate

To further understand the computations leading to the monkey’s aborting behavior we trained RL agents instantiating different neural architectures to navigate to the location of hidden targets, and examine if and how they abort offers (**Fig. 2A**). We formulate the “firefly task” ^20, 21, 24–26^ as a Partially Observable Markov Decision Process (POMDP^30, 31^). Specifically, at each time step *t*, the state of the environment ***s***_*t*_includes the agent and target position, as well as the agent’s velocity. The agent then makes an observation given the state, **o**_*t*_ ∼ *O*(**o**_*t*_|***s***_*t*_), and takes an action **a**_*t*_ controlling its linear and angular velocity. The action updates the state to ***s***_*t*&1_, according to the environment transition probabilities *T*(***s***_*t*&1_|***s***_*t*_, **a**_*t*_), i.e., the task dynamics. The agent receives reward *r*_*t*_ determined by the reward function *R*(***s***_*t*_, **a**_*t*_), which is a positive scalar if the agent stops within the reward zone (65 cm, as for the macaques), and 0 otherwise.

**Figure 2.**
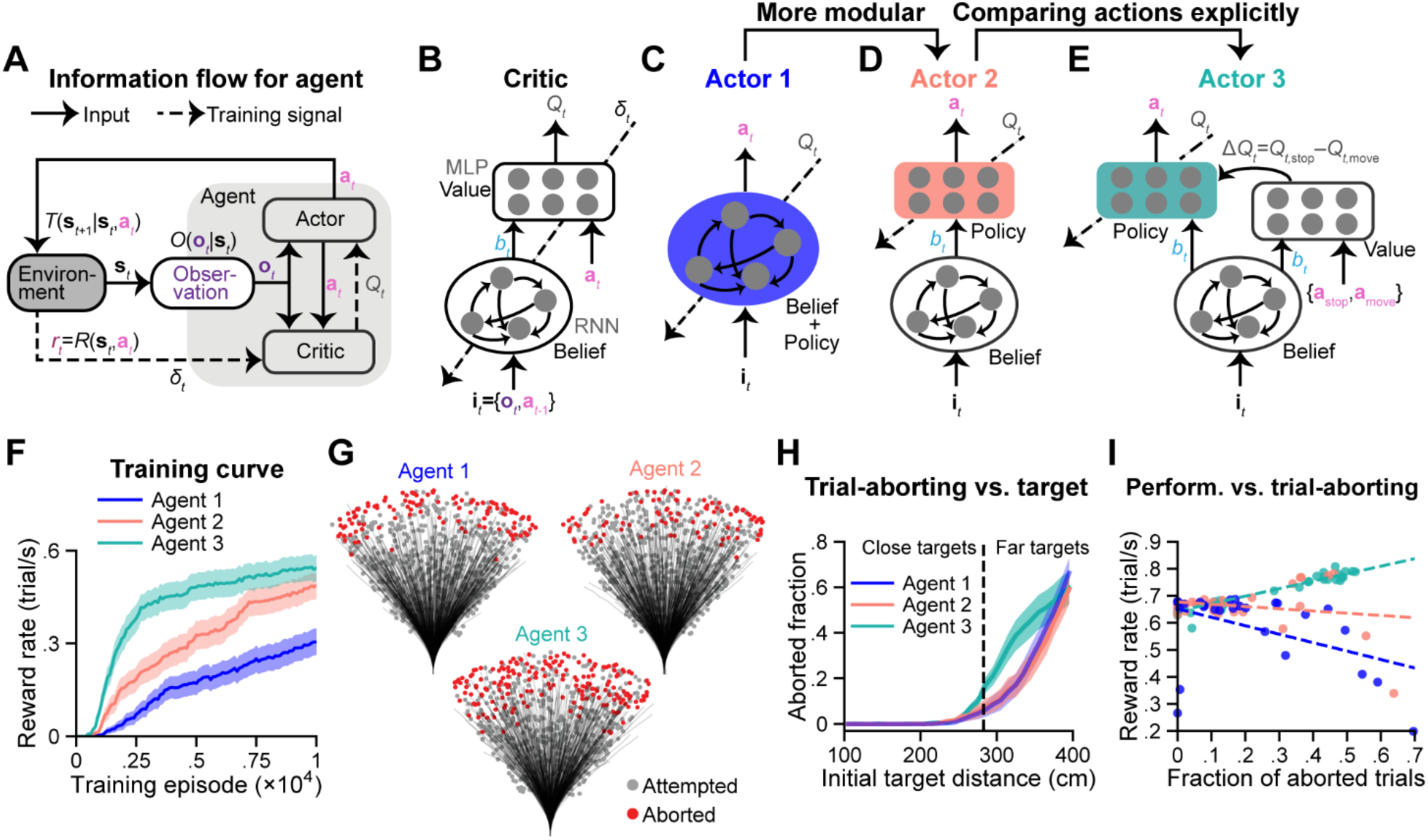
Modular RL actors augmented with specific model-free critic values strategically abort trials to maximize reward rate. **A.** Block diagram showing the interaction between an RL agent and the task environment. At each time step *t*, the agent receives a partial and noisy observation **o**_*t*_ of the state ***s***_*t*_. The actor network integrates the observation with past observations (via the current belief) and then outputs an action **a**_*t*_, which partially updates ***s***_*t*_to ***s***_*t*"1_(the state also has its own dynamics). The actor is trained to output the action that maximizes the value *Q*_*t*_ according to the critic, while the critic is trained to generate more accurate value estimations using the TD error δ_*t*_after receiving reward *r*_*t*_from the environment (Eq. 2). **B.** Schematic of critic. A critic network comprises two modules. The belief module is implemented as an RNN, integrating available information **i**_*t*_=(**o**_*t*_,**a**_*t*$1_) to compute a belief ***b***_***t***_about the state. The value module outputs the value *Q*_*t*_of an action **a**_*t*_given ***b***_***t***_. Both modules are trained end-to-end through back propagation. **C.** Schematic of Actor 1 implemented with a single RNN. Given **i**_*t*_, Actor 1 needs to compute both belief and action together. **D.** Schematic of Actor 2, which uses the belief module trained by the critic in (**B**) to construct *b*_*t*_. Then, a policy module computes an action **a**_*t*_ from its input *b*_*t*_. Only the parameters in the policy module are updated in actor training. **E.** Schematic of Actor 3. Similar to (**D**), this network uses the belief module from (**B**) to construct *b*_*t*_. However, it also directly receives Δ*Q*_*t*_ = *Q*_*t*,s()*_ − *Q*_*t*,+)ve_ from the critic MLP. Actor 3 takes an action **a**_*t*_ given both *b*_*t*_ and Δ*Q*_*t*_. **F.** Reward rate (rewarded trials per second) during training, measured using a validation set (500 trials) for each agent at each checkpoint (every 100 training episodes). Shaded regions denote ±1 s.e.m. across 50 training runs with different random seeds. **G.** Overhead view of the spatial distribution of 800 representative targets and trajectories for Agents 1, 2, and 3 (sampled from random seeds that yielded reward rates > 0.2) navigating toward these targets. Red/gray dots denote aborted/attempted targets, as Fig. 1. **H.** Fraction of aborted trials versus initial target distance, averaged across random seeds. Shaded regions denote ±1 s.e.m. For each seed, target distance is binned for every 10 cm, and the aborted fraction is calculated in each bin. The black dashed line denotes the median distance (200√2 cm) of the targets. **I.** Reward rate versus aborted fraction for 2000 targets. Each dot denotes a training run with a unique random seed. Dashed lines denote the linear regression lines for the dots.

To generate behaviors, we adopt an actor-critic approach^3, 32, 33^ in training artificial agent to perform this task (**Fig. 2A**). At each time *t* the actor aims to generate an action **a**_*t*_that maximizes the state-action value, which represents the expected sum of discounted rewards from time *t* onward:

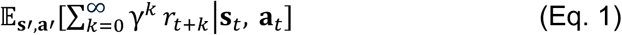

where γ is the temporal discount factor and ***s***′ and **a**′ denote future states and future actions. Since there is no terminal state after a trial concludes, the selection of actions considers not only the reward associated with stopping at the target location in the current trial, but also the potential rewards for all future targets. This differs from our previous work^22^, in which the state-action value function was bound to the current trial (i.e., there was no consideration of time-frames outside the current trial). A critic network (**Fig. 2A, B**) is used to approximate state-action values, denoted as *Q*_*t*_, given ***s***_*t*_and **a**_*t*_. The critic’s approximation is updated using the temporal-difference reward prediction error^34^ (TD error) δ_*t*_,

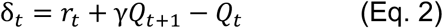

where *Q*_*t*&1_ is the next-step value obtained through bootstrapping (for details see *Methods* and *Eqs. 1 8, 19*).

In our task, observations **o**_*t*_ only contain partial information about the state. For instance, the target is only visible for a short duration (i.e., the 300ms over which it is visible). Likewise, self-position of the agent has to be inferred from velocity observations (i.e., “path integration”) and is not directly observable (i.e., there is no landmark information in this task). Hence, the critic network first needs to construct an internal state representation, or ‘belief’ *b*_*t*_, about the current state (the actual state being a feature the agent does not have access to) in order to approximate belief-action values. This belief is computed via a recurrent neural network (RNN; **Fig. 2B**) that enables the temporal integration of evidence and retains this information in its activity state. The input, **i**_*t*_, for the RNN includes the noisy observation **o**_*t*_ (self-velocities, and target position when the target is visible) and the previous action **a**_*t*"1_ as an efference copy. Next, the critic approximates *Q*_*t*_ given *b*_*t*_ from the RNN and an action **a**_*t*_. This computation is accomplished via a feedforward (i.e., memory-less) multi-layer perceptron (MLP; **Fig. 2B**, see^22^ for the advantages of this modular architecture). Note that the critic including the RNN and MLP is trained end-to-end through back-propagation with the objective of learning *Q*_*t*_. Nevertheless, in Zhang et al.^22^ we showed that even without an explicit objective to shape *b*_*t*_, the learned belief is near-optimal, and the RNN and MLP are respectively specialized in computing belief and value, without intermixing the computations of these variables.

Based on its belief, an actor network is trained to select an action **a**_*t*_that maximizes the value *Q*_*t*_according to the critic. All of our agents use the same modular critic architecture, while we explore three different actor architectures. Actor 1 (**Fig. 2C**) has the simplest: a holistic architecture. Given the state-related inputs **i**_*t*_, it uses a single RNN to compute both belief and action, thus lacking architecture-enforced neural specializations. Actor 2 (**Fig. 2D**) and Actor 3 (**Fig. 2E**) enforce neural modularity by providing separate modules that have structures naturally suited to computing each variable. Both actors inherit the critic’s RNN to compute *b*_*t*_, and then generate **a**_*t*_using an MLP that is optimized to maximize *Q*_*t*_. Actor 3 is further granted an additional input: the difference in value Δ*Q*_*t*_ (evaluated by the critic’s MLP) between stopping (**a**_s/01_) vs. executing a movement and thus continuing the trial (**a**_2034_):

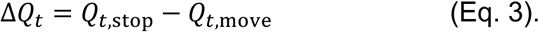

In other words, the policy module in Actor 3 (**Fig. 2E**, green) is not only influenced by *b*_*t*_from the critic’s belief RNN, but also by an explicit value signal from the critic’s value module. In this sense, the critic’s role in actor learning is two-fold. For all actors, it serves as the learning objective to adjust the actor’s weights (a slow process^9^). For Actor 3, it additionally serves as an internal model capable of evaluating actions of interest, such as **a**_s/01_ and **a**_2034_, to influence policy neurons’ dynamics (a fast process^9^, see *Discussion* for more detail).

Among these agents, Agent 3 exhibited the fastest learning rate and the highest final performance (**Figs. 2F, Fig. S3A**). It also had the most training instances (i.e., random seeds) achieving proper learning and showing the development of aborting behavior (**Fig. S3B, C, D**). Indeed, the behavior of these agents showed that while all networks could abort trials (see **Fig. 2G** for example sessions), this was more frequent in Agent 3 than Agents 1 or 2 (**Fig. S3D,** K-S test: Agents 1 vs. 3, *p* = 0.018; Agents 2 vs. 3, *p* = 0.002). Furthermore, the spatial distribution of aborted trials differed: Agent 3 aborted trials for closer distances than the other agents (**Fig. 2H, S3E**). Moreover, aborting was only beneficial for Agent 3: reward rate increased with trial-aborting behavior in Agent 3 (**Fig. 2I**: Pearson’s *r* = 0.95, *p* = 5.45 × 10^"21^; slope= 0.28), but not for Agents 1 or 2. Indeed, aborting trials hindered performance in Agent 1 (*r* = −0.44, *p* = 0.023; slope= −0.31) and did not enhance performance in Agent 2 (*r* = −0.2, *p* = 0.24). This suggests that while the aborting behavior was adaptive for the former, it was not for the latter two. Interestingly, when we double the training of Agent 1 (**Fig. S3F, G, H, I**), the negative correlation between aborting behavior and reward rate became less significant (**Fig. S3J**: *r* = −0.22, *p* = 0.2; slope= −0.14), demonstrating that while aborting may ultimately be beneficial for in Agent 1, it develops much earlier in Agent 3.

Together, these results suggest that strategically aborting trials to increase reward rates (1) requires evaluating a value function that accounts for long-term value beyond just a single trial (which our past work neglected^22^), (2) benefits from a modular actor architecture (Actor 2 and 3 vs. 1; also see **Fig. S4** and neural specialization analyses in *Methods*), and (3) can be catalyzed by providing actor networks the difference in value between performing the task and not performing it (Actor 2 vs. Actor 3).

### Value-informed policy extraction explains reward-maximizing trial-aborting behavior

Next, we aimed to understand the mechanism by which Agent 3 uses its additional learning signal to increase its reward rate by aborting trials. While poor RL performance is often attributed to an imperfect value function^35, 36^, here Actors 2 and 3 maximize the same type of value functions (their respective critics) and are provided with the same type of beliefs (from critics). The difference between these agents, therefore, ought to be in their actors’ abilities to extract an optimized policy from the critics (see^37^ for a similar argument). To demonstrate this, we compute Δ*Q*_*t*_ (Eq. 3) for Agents 2 and 3 (see ‘Critic’s opinion’ section in *Methods*). This difference in value is provided as a learning signal to Agent 3, but can also be computed for Agent 2, as any critic can estimate values for the “move” and “non-move” (i.e., stop) actions. **Figure 3A** shows that while the critics of Agents 2 and 3 have similar “opinions” about the value, Δ*Q*_*t*,1_, of aborting (red) or attempting (gray) trials when target is presented (Agent 2: top, vermilion; Agent 3: bottom, green), the actor’s behavior in Agent 3 followed its critic’s opinion more closely. This is further illustrated by contingency tables (**Fig. 3B**) and the fraction of trials in which the actor attempted a trial despite the critic’s suggestion to abort (**Fig. 3C**, Agent 2: 46%, Agent 3: 17%).

**Figure 3.**
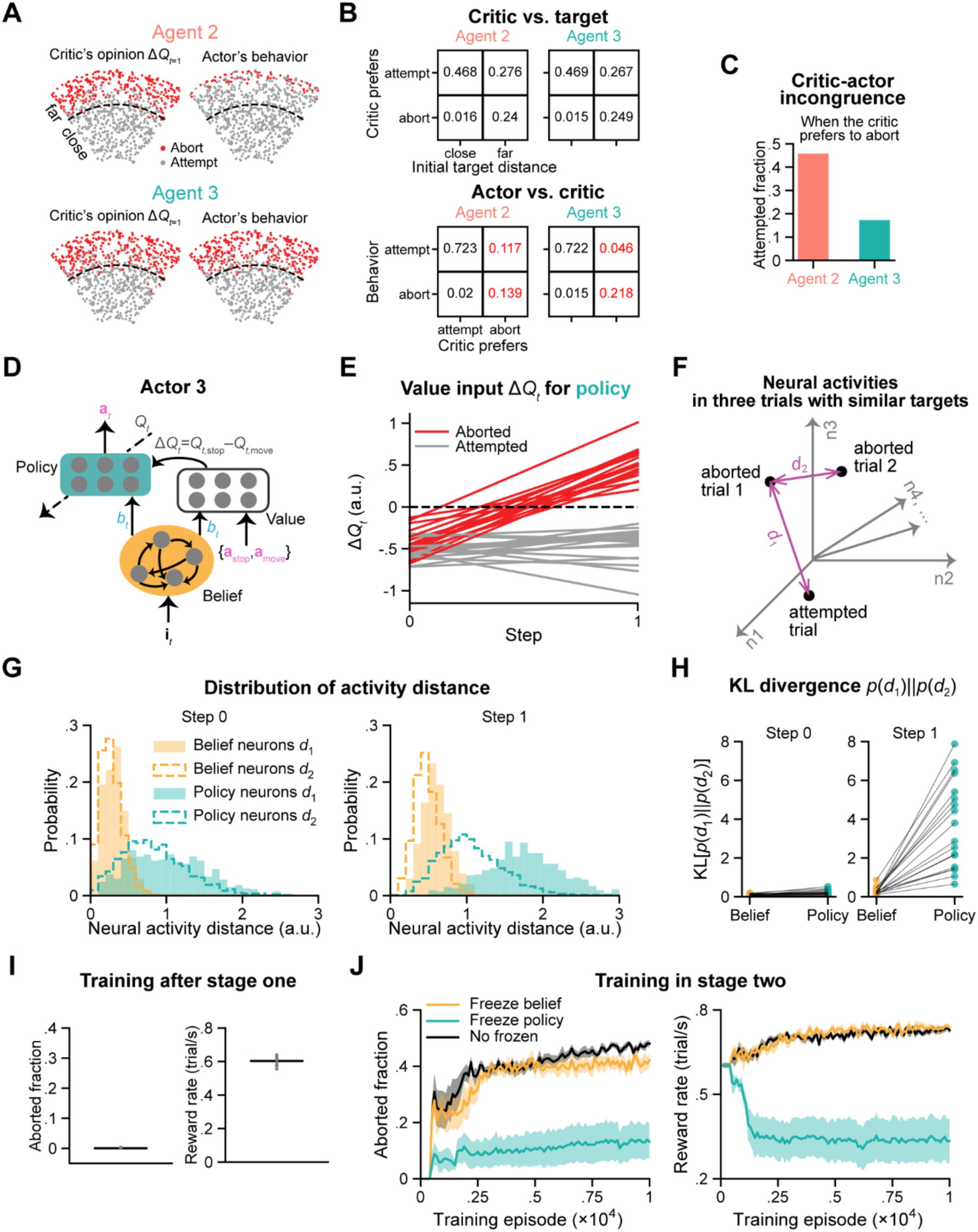
Policy neurons reflect a trial-aborting strategy while belief neurons do not. **A.** Overhead view of the spatial distribution of 800 representative targets for a training run of Agent 2 (top) or Agent 3 (bottom). Red/gray dots on the left denote that agents’ critics valued abort/attempt more (Δ*Q*_*t*.1_>0 / Δ*Q*_*t*.1_<0), while on the right they denote whether trials were aborted/attempted based on behavior. Black dashed arcs equally separate the target sampling area by target distance. **B.** Numbers in the tables denote fractions of total trials. *Top*: Contingency tables showing the critic’s preference (Δ*Q*_*t*.1_) vs. initial target distance. *Bottom*: Table showing the actor’s behavior vs. the critic’s preference. Red color highlights pronounced differences between agents 2 and 3. Trials from all random seeds (with reward rate > 0.2) were included. **C.** Fraction of attempted trials (the red number in the first row of each table in **B**) among trials that the critic preferred to abort (column sum of red numbers in each table in **B**) for two agents. **D.** Schematic of Actor 3, shown as in Fig. 2E. We use yellow to denote beliefs, both for the module in this panel and for traces in subsequent panels. **E.** Δ*Q*_*t*_ computed by the value module in **D** during the first two timesteps of trials. 2000 trials were used to evaluate the agent with each random seed. Each line denotes Δ*Q*_*t*_ either averaged across aborted trials (red) or across target-matched attempted trials (gray) for each seed. **F.** Illustration of measuring distances in high-dimensional neural activity space between activities for an aborted and a target-matched attempted trial (*d*_1_), or between activities for two target-matched aborted trials (*d*_/_). **G.** Distributions of *d*_1_ and *d*_/_ at timestep 0 (left) and timestep 1 (right) for belief and policy neurons in an agent trained with a random seed. **H.** KL divergence between distributions of *d*_1_ and *d*_/_ in **G**, including all random seeds (each dot denotes a random seed). **I.** Aborted fraction and reward rate after training stage one, averaged across training instances. 500 trials were used. Gray dots: 10 training instances. **J.** Fraction of aborted trials (left) and reward rate (right) over stage two training. After training stage one (I), the agent then continued to train with either belief or policy neurons frozen, or with none of the neurons frozen. 500 trials were evaluated for each agent at each checkpoint, occurring every 100 training episodes. Shaded regions denote ±1 s.e.m. across training instances.

To further understand the mechanism by which policies may be extracted for a given “opinion”, we may also examine the behavior of populations of artificial neurons within Agent 3’s actor—the network that appropriately aborts trials to maximize the reward rate. This actor contains a belief module (**Fig. 3D**, orange) and a policy module (**Fig. 3D**, green). At trial onset, Δ*Q*_*t*,-_ is matched (and negative) for subsequently aborted (red, **Fig. 3E**) and attempted (gray, **Fig. 3E**) trials. In other words, when the target is first presented, the value of moving is higher than the value of stopping (Δ*Q*_*t*,-_ is negative). This is because to start (and then potentially abort) a trial, the agents first have to move. If they do not move, the trial will last until a time-out, akin to the monkey’s “never start” trials (see ‘Reward’ section in *Methods*). Immediately after the initial step, however, Δ*Q*_*t*,1_ becomes positive for subsequently aborted (*Q*_*t*,s()*_ > *Q*_*t*,+)ve_) trials but not for attempted ones. Thus, we may reason that if units were reflecting Δ*Q*_*t*_, there should be a divergence in neural activity for subsequently aborted vs. attempted trials already at the second time-point after targets are presented.

To examine this hypothesis, in Agent 3 we matched each aborted trial with (1) its most similar attempted trial (in terms of targets’ Euclidean distance upon presentation) and (2) its most similar aborted trial. Then, we expressed the population activity in a high-dimensional space (**Fig. 3F**) and compared the Euclidian distance between aborted and attempted trials (*d*_1_), as well as between two aborted trials (*d*_2_) at trial onset (*t* = 0) and target presentation (*t* = 1) in the belief (orange) and policy (green) modules. We repeated this procedure for all aborted trials in a 2000-trial session to estimate full distributions of these distances (**Fig. 3G**). Results showed that, at trial onset, belief and policy neurons do not differentiate between attempted and aborted trials more than they do between pairs of aborted trials (**Fig. 3G**, left). In contrast, after the initial step, the neural state difference between attempted and aborted trials (*d*_1_) is distinct from that between two aborted trials (*d*_2_) within the policy module, but not in the belief module (**Fig. 3G**, right). This effect can be quantitatively assessed by calculating the KL divergence between the distributions of *d*_1_ and *d*_2_ (**Fig. 3H**).

To further verify the role of the belief and policy modules, we conducted ablation analyses by freezing neurons during the learning of trial-aborting behavior (see ‘Agent Training’ section in *Methods*). We first trained agents to navigate to targets without considering long-term outcomes, i.e., only considering the target in the current trial (training stage one). Namely, agents attempted to maximize

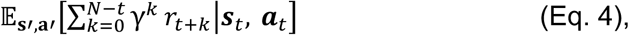

where *N* denotes the final step in a trial. In this stage, agents did not abort trials (**Fig. 3I**). Then, in stage two, we reverted the definition of value to that in *Eq. 1* by changing the upper bound of the sum from *N*–*t* to ∞: we now considering not only the current target up to time *N*, but also all times for all subsequent targets. We continued training the agent while either (1) freezing parameters in the belief module (**Fig. 3D**, orange), (2) freezing parameters in the policy module (**Fig. 3D**, green), or (3) allowing all parameters to be updated. The results show that preventing the belief module from learning during the second stage of training impacted neither the agent’s trial-aborting behaviors (**Fig. 3J**, left, orange) nor performance (**Fig. 3J**, right, orange) compared to the agent with no frozen parameters (**Fig. 3J**, black). In contrast, freezing the policy module (**Fig. 3J**, green) during stage two training resulted in very poor behavior.

Together, these results suggest that reward-enhancing aborting behavior is supported by a modular actor network whose policy module is conditioned on a specific value learning signal allowing for better (or faster) alignment with the critic. Further, this reward enhancing behavior is supported by the policy module but not the belief module of the actor, as evidenced by artificial neural activity at target presentation and confirmed through ablation (neuron-freezing) analyses.

### Dorsolateral prefrontal cortex as a value-informed policy network optimizing session-long reward rates

The RL modeling suggests that session-long aborting behaviors may be learned faster by a modular policy module directly informed of value differentials between performing the task at hand or not. This difference is evident in the agent’s neural activity as soon as target presentation. We next investigated whether this pattern occurs in real neural activity of macaques performing the “firefly task” ^20, 21, 24–26^.

We recorded single-unit spiking activity from the dorsolateral prefrontal cortex (dlPFC; 445 single units), parietal area 7a (2647 single units), and the dorso-medial superior temporal area (MSTd; 240 single units). In prior work we have solely analyzed “attempted trials” (**Fig. 1B**, leftmost), and established that coding of the key latent variable within this task (i.e., the instantaneous and unobservable distance to target) is strikingly distributed (e.g., is decodable from dlPFC, 7a, and MSTd^25^) and affected by recurrence, involving bi-directional coupling between sensory and cognitive areas^24^. This distributed and recurrent nature of the coding of the key latent variable (i.e., “belief”) is akin to the RNN computing beliefs in our RL agents. Here, instead, we aim to establish whether we can identify biological units matching the properties of the agent’s policy module (e.g., MLP in Actor 3, **Fig. 2** and **3**).

As for the artificial units in Actor 3, we matched each aborted trial to its most similar attempted trial in terms of target distance, eccentricity, and afforded reward rate. Further, we only examined neural responses during target presentation (−0.3 to 1 second post-stimulus onset): within this window, motor output is not yet different between attempted and aborted trials. Indeed, **Figure 4A** shows the subsampled distribution of matched target locations for attempted (gray) and aborted trials (red; top row) for three example sessions, as well as their velocity profiles (second row) within the time period of interest (see **Fig. S5** for additional example sessions). Next, we constructed raster plots (**Fig. 4A**, third and fourth row) and peri-stimulus time histograms (PSTHs; **Fig. 4A**, fifth row) for neurons in each region, based on this subsampled set of trials. We found neurons in MSTd (**Fig. 4A**, leftmost), area 7a (**Fig. 4A**, middle), and dlPFC (**Fig. 4A**, rightmost) that seemingly differentiated between attempted and aborted trials (Wilcoxon sign-rank test, baseline-corrected spike counts between 0 and 0.3 seconds after stimulus onset; *p* < 0.05). Responses from these neurons were heterogenous, but importantly, they did not merely show a lack of response during aborted trials, as if the animals had ‘missed’ the target presentation. On the contrary, on occasion (e.g., **Fig. 4A**, dlPFC example) the response to aborted trials was greater than that to attempted ones. Further, the vast majority of neurons that differentiated between subsequently aborted or attempted trials did so within the first 300ms following target (i.e., offer) presentation.

**Figure 4.**
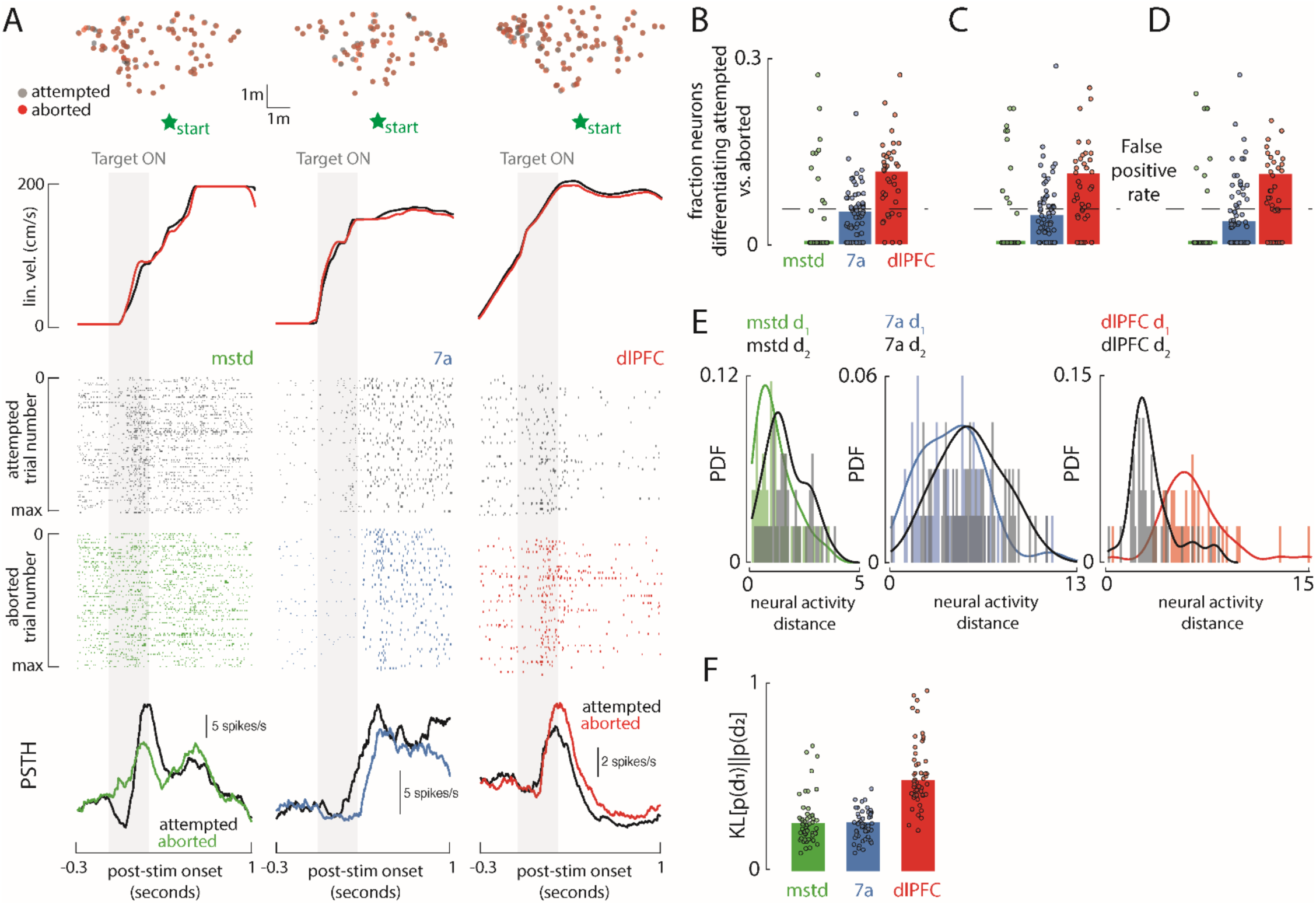
Dorsolateral prefrontal cortex reflects differing policies. **A.** Data from 3 example cells, one each from MSTd (green; 1^st^ column), area 7a (blue; 2^nd^ column), and dlPFC (red; 3^rd^). Top row shows the distribution of matched target locations (overhead view) for attempted (gray) and aborted (red) trials. Starting location of the animal is shown by a green star. The second row shows the average linear velocity profile for attempted (gray) and aborted (red) trials. This demonstrates that in the period surrounding target presentation (-0.3 to 1 second post-stimulus onset), the velocity profiles were matched across trial types. The transparent gray area shows the period of time the target was visible (0 to 300ms after stimulus onset). Third and fourth row respectively show example raster plots for attempted (black) and aborted trials (colored as a function of brain area). Each row is a trial and each dot is a spike. Fifth row shows the peri-stimulus time histogram (PSTH) of the data presented in the raster plots (aborted in black and attempted as a function of brain area). **B.** Fraction of neurons differentiating in their peri-stimulus time response (spike count from 0 to 300ms after stimulus onset, baseline corrected) between attempted and aborted trials. Each dot is the fraction computed on individuals sessions. Green (MSTd), blue (7a), and red (dlPFC) bars are the median across sessions. Data from MSTd (due to its lower total yield) is not Gaussian so we performed non-parametric significance testing (Mann-Whitney U test). Dashed black line shows the false positive rate at 5%, given the significance threshold employed. **C.** Same as **B.,** when only considering trials where the animal was stopped (linear velocity = 0 cm/s and angular velocity = 0 deg/s) before stimulus onset (–300 to 0 ms). **D.** Same as **C.,** when additionally filtering out trials wherein the animal aborted within 1 trial of stimulus onset. **E.** Probability density functions of *d*_1_ (colored; Euclidean distance in high-dimensional space between matched attempted and aborted trials) and *d*_/_ (black; distance between matched aborted trials) in MSTd (green), area 7a (blue), and dlPFC (red). **F.** KL divergence between distributions shown in **E.** For statistical testing, we subsampled (without replacement) 50 times (dots) from each distribution, for 20 sessions. Bars shows the means across subsamples.

A greater fraction of neurons in dlPFC differentiated between policies (i.e., attempt vs. abort) for a matched offer compared to the other areas. In fact, 12.6% of neurons in dlPFC (**Fig. 4B**, red, median across session) distinguished between attempted and aborted trials (baseline-corrected spike counts between 0 and 0.3 seconds post-stim onset; p<0.05). This fraction is greater than that observed in area 7a (4.3%; Mann-Whitney U, p = 7.8×10^-5^) and MSTd (0% median; p = 7.7×10^-7^), and significantly above chance (p = 0.02, by definition 5%). In contrast, area 7a (p = 0.53) and MSTd (p = 1.0) did not have more neurons than expected by chance that were sensitive to different policies during target presentation (Mann-Whitney U test). These observations remained true when we required animals to be immobile before target presentation (i.e., linear speed = 0cm/s; **Fig. 4C**; dlPFC fraction vs. chance, p = 0.018). Likewise, the same results held when additionally filtering for trials wherein aborting did not occur for at least 1 second after stimulus offset (as in **Fig. 4A**, i.e., matching pre- and post- stimulus velocity profiles; **Fig. 4D**, dlPFC fraction vs. chance, p = 0.007).

To examine population-level responses (as opposed to a series of individual neurons) we performed the same analysis as for the model in **Fig. 3F-H**. That is, simultaneously recorded neural activity from a given area defined a high-dimensional space (average firing rate 0 to 300ms post-stimulus onset), with each neuron representing an independent axis. Then, we paired each aborted trial with two other trials from the same session that had the most similar targets, one attempted and one also aborted. Then, we computed the Euclidian distances within these high-dimensional spaces, and contrast the distance in neural activity between pairs of aborted-attempted trials (*d*_1_) or pairs of aborted trials (*d*_2_) for each brain region (akin to **Fig. 3F**). As shown in **Figure 4E**, the results demonstrate a separation in neural space between attempted and aborted trials that exceeds that for paired aborted trials in dlPFC (red), but not 7a (blue) or MSTd (green). This effect can be quantified by calculating the KL divergence between distributions of *d*_1_ and *d*_2_ for each brain region (**Fig. 4F**). Examining the component of neural activity along the top principal component within either dlPFC, 7a, or MSTd, we corroborated that only dlPFC distinguishes between attempted and aborted trials upon target presentation (see **Fig. S6**).

## Discussion

We developed a task wherein animals naturalistically forage for reward in virtual reality^19–22, 24–26, 42^. A central feature of this task, just as in real-world foraging, is the implicit ability and importance of *manipulating* (i.e., minimizing) the time between *possible* rewards. This breaks with the mold of traditional systems neuroscience work which, by and large, experimentally determines the speed at which an agent can acquire rewards (e.g., the speed of stimulus presentation and the duration of inter-trial intervals). Similarly, in traditional approaches, animals usually have a very limited ability to impact the structure of their tasks: for example, how valuable the probable next offer is relative to the current (see^43–49^ for recent exceptions using ‘time’ as a key experimental variable to estimate value^43, 44^ or confidence^45–49^). Instead, in our task, animals can abort trials, saving themselves up to tens of seconds. This behavior, however, requires forgoing the potential for reward on the current trial.

Using this task, here we demonstrate (1) that macaques have knowledge about their own performance and strategically use this knowledge in aborting trials in a session- and individual-specific manner to increase session-long reward rates, (2) that computationally this behavior requires considering state-action values beyond a single trial, and (3) that this behavior is learned faster with a modular actor-critic architecture in which a policy module is given an explicit estimate of the relative value of aborting the current offer. This explicit signal makes it easier for the actor to use the critic’s evaluations (*policy extraction*) within the actor-critic architecture. Finally, we show that (4) individual neurons and (5) population dynamics in dlPFC but not parietal area 7a or MSTd reflect the strategic aborting of trials upon target presentation—even if the aborting itself won’t occur for another second or longer. The fact that a putative policy module (i.e., “dlPFC”) reflects subsequent aborting behavior already during target presentation is as predicted by the modular actor-critic artificial neural network.

Previously, we have shown that coding of distance to target—the key latent variable that must be minimized in this task—is distrusted and recurrent^24, 25^. That is, single units across cognitive and sensory cortices (i.e., dlPFC, 7a, and MSTd) are tuned to the instantaneous angular and/or radial distance to target^24^. Dorsolateral prefrontal cortex and MSTd appear to have a preference for distal targets, and preferentially couple with one another when animals keep track of the (invisible) target position with their gaze^24^ (i.e., an embodied strategy). Area 7a is more frequently active as targets get closer^24^ and is the area most clearly encoding sensorimotor state variables (e.g., linear velocity). Likewise, distance to target is decodable from all three of these areas^25^: it is more invariant to sensory context in dlPFC and 7a, and is more modulated by observations in MSTd. Together, these previous results suggest that MSTd and 7a are strongly modulated by observations and state variables (also see^26, 50^), while the encoding of beliefs are more distributed. Interestingly, the current study is the first time we see clear area specialization within this “firefly task”: dlPFC but not other areas reflect the animal’s policy.

More broadly, the results add to our understanding of reinforcement learning within fronto-striatal circuits, and evoke a number of questions and predictions. For instance, most neuroscience tasks typically involve learning state values, and the ventral striatum is considered to play a key role in this process^51–53^. However, in more complex tasks, value depends on both state and actions. Previous studies suggest that both action values and state-action values are computed in the dorsal striatum (DS), which may evaluate prospective actions represented in the dlPFC^53–55^. Interpreting the computations of our favored model, Agent 3, we suggest that the value module in the critic mimics the DS, computing values for actions proposed by the policy module in the actor, the dlPFC (see **Fig. S7A**). Neurobiologically, these interactions would need to be mediated by other areas, because the dorsal striatum does not directly project to dlPFC. Instead, it projects to the dorsal pallidum, which then projects back to medial-dorsal nucleus of the thalamus, and finally back to dlPFC (**Fig. S7B**). Thus, these relay areas between dorsal striatum (i.e., the “critic”) and dorsolateral prefrontal cortex (i.e., the “actor”) ought to also be informed of the value of performing specific state-actions pairs.

We also show that allowing the value module to project values for “stopping” and “moving” actions to the policy module improves learning rates and task performance by allowing animals to abort low reward rate offers. Importantly, this added signal to the policy module does not change value computations, but allows for an optimized policy extraction (i.e., critic and actor becoming more congruent with one another). In this same vein, prior studies have shown that striatal medium spiny neurons expressing D1 or D2 dopamine receptors, which respectively facilitate or inhibit actions, correlate positively and negatively with state-action values^56, 57^. These results support the argument that D1 and D2 striatal neurons first decide whether or not to initiate an action, with specific actions being selected thereafter^58^. Our results align with this view, suggesting that the dlPFC may modulate the balance between D1 and D2 neurons in deciding whether or not to engage with a particular offer. Seemingly, this two-step approach (first decide whether to act, then decide on the particular action), optimizes policy extraction. This interpretation is also in line with hierarchical RL and the suggestion that dlPFC may play a particularly important role in the arbitration between different and potentially concurrent RL algorithms that may exist in biological networks^59–62^.

To further test the putative role of dlPFC as a value-informed policy module, in future work it will be important to test dlPFC dynamics during learning of the firefly task, or another task (1) engaging model-based RL^21, 22^, (2) allowing for different policies given a matched offer, and (3) indexing naturalistic closed-loop behaviors wherein time-to-reward is manipulable. Moreover, we may test multiple state-transition dynamics (e.g., target distances that are correlated across trials) and simultaneously record from candidate striatal and cortical actor and critic modules. Likewise, it will be informative to conduct causal studies such as lesioning or electrical stimulation. Lastly, given that the central benefit of meta-RL is the almost instantaneous tackling of novel scenarios, it will be crucial in future work to consider the neural correlates and algorithms subserving multiple tasks, and to investigate how dlPFC allows for this generalization.

In conclusion, we demonstrate that macaques may reason about their own long-term performance and idiosyncratically decide what experimental offers are worth working for, and which are not. Their aborting of trials is not random, but instead acts to increase reward rates (i.e., they abort low reward rate offers). This computation is best recapitulated by a modular actor-critic architecture in which a policy module is explicitly informed of the relative value of performing the task, or not. This additional signal improves the use of value estimates, and leads the policy module to distinguish between subsequently aborted and attempted trials already during offer (i.e., target) presentation. We observe a similar behavior for neurons in dlPFC, but not in 7a or MSTd in macaques.

## Methods

### Animals, Animal Shaping, and Dataset

Three adult male rhesus macaques (*Macaca Mulatta*; 9.8-13.1kg) were studied. We analyzed behavioral and neural data from 30 sessions in monkey S, 28 sessions in monkey Q, and 14 sessions in monkey M, resulting in a total of 72 sessions. The animals performed an average of 1355 trials per session, resulting in a total dataset of 97591 trials. These trials/sessions have been reported on before^24, 25^, yet in those cases we focused exclusively on attempted trials and disregarded failure modes: aborted trials, “never started” trials, and “never stopped” trials. Here, our focus is in understanding the neural underpinning of aborted vs. attempted trials, which constituted an independent dataset from those previously reported.

Animals were chronically implanted with a lightweight polyacetal ring for head restraint. They were then trained via standard operant conditioning to path integrate to the location of briefly visible targets (“fireflies”) by accumulating optic flow velocity signals^42^. All surgeries and procedures were approved by the Institutional Animal Care and Use Committee at Baylor College of Medicine and New York University, and were in accordance with National Institutes of Health guidelines.

### Experimental Setup

Monkeys were head-fixed and secured to a primate chair. To navigate, the animals used an analog joystick (M20U9T-N82, CTI electronics) with two degrees of freedom controlling their linear and angular speed in a virtual environment (linear max = 200cm/s; angular max = 90deg/s). A DLP projector (3-chip Christie Digital Mirage 2000, Cypress, CA, USA) rear-projected images onto a 60 x 60cm screen that was attached to the front of the field coil frame, 32.5cm from the monkey. The projector rendered stereoscopic images generated and rendered using C++ Open Graphics Library (OpenGL; Nvidia Quadro FX 3000G) by continuously repositioning a virtual camera (60Hz; height of 1m) based on the monkey’s joystick inputs. The virtual environment was composed of ground plane textural elements with a lifetime of 250ms (at exception for their first presentation, in which each element had a different lifetime drawn from a uniform distribution between 0 and 250ms). These elements were isosceles triangles (base x height: 8.5 x 18.5 cm^2^) that were randomly repositioned and reoriented at the end of their lifetime. The density of the ground plane elements was 5.0 elements/m^2^. The ground plane was circular, with a radius of 70m (near and far clipping planes at 5cm and 40m, respectively). The animals were re-positioned at the center of this environment at the beginning of each trial. Importantly, this teleportation was undetectable to animals, given that we matched the last frame of trial *t-1* and first of trial *t*. In other words, from the standpoint of the animals, they continuously explored an infinite environment. Spike2 software (Cambridge Electronic Design Ltd., Cambridge, UK) was used to record and store the timeseries of the target and animal’s location, animal linear and angular velocity, as well as eye movements. All behavioral data were recorded along with the neural event markers at a sampling rate of 833.33Hz.

### Behavioral Task

Monkeys steered to a target location (circular disc of radius 20 cm) that was cued briefly (300ms) at the beginning of each trial. Each trial was programmed to start after a variable random delay (truncated exponential distribution, range: 0.2 – 2.0s; mean: 0.5s) following the end of the previous trial. The targets appeared at a random location between –45 to 45deg of visual angle, and between 1 and 4m relative to where the animal was stationed at the beginning of the trial. The joystick was always active, and thus monkeys were free to start moving before the target vanished, or before it appeared. Monkeys typically performed blocks of 500-750 trials before being given a short break. Feedback was given in the form of juice reward following a variable waiting period (truncated exponential distribution, range: 0.1 – 0.6 s; mean: 0.25 s) after stopping within 0.6m from the center of the target (“reward boundary”). No juice was provided otherwise.

### Neural Recording and Pre-Processing

We recorded neural activity extracellularly, either acutely using a 24-channel linear electrode array (100 µm spacing between electrodes; U-Probe, Plexon Inc, Dallas, TX, USA) or chronically with multi-electrode arrays of either 48 or 96 electrodes (Blackrock Microsystems, Salt Lake City, UT, USA). Parietal area 7a and dorso-medial pre-frontal cortex (dlPFC) were recorded both with linear probes and Utah arrays, while dorso-medial superior temporal area (MSTd), being a deep structure, was only recorded with linear arrays. We confirmed that our results do not depend on type of electrode used (also see Noel et al., 2022). During acute recordings with the linear arrays, the probes were advanced into the cortex through a guide-tube using a hydraulic micro-drive. Spike detection thresholds were manually adjusted separately for each channel to facilitate real-time monitoring of action potential waveforms. Recordings began once waveforms were stable. The broadband signals were amplified and digitized at 20 KHz using a multichannel data acquisition system (Plexon Inc, Dallas, TX, USA). Raw data was stored along with the action potential waveforms for offline analysis. Additionally, for each channel, we also stored low-pass filtered (-3dB at 250Hz) local-field potentials (LFP). For the array recordings, broadband neural signals were amplified and digitized at 30 KHz using a digital headstage (Cereplex E, Blackrock Microsystems, Salt Lake City, UT, USA), and stored for offline analysis. Additionally, for each channel, we stored low-pass filtered LFPs sampled at 500Hz. Finally, copies of event markers were received online from the stimulus acquisition software (Spike2) and saved alongside the neural data.

Spike detection and sorting were performed on raw neural signals using KiloSort 2.0 on an NVIDIA Quadro P5000 GPU. The spike clusters produced by KiloSort were visualized in Phy2 and manually refined by a human observer by either accepting or rejecting KiloSort’s label for each unit. In addition, we computed three isolation quality metrics; inter-spike interval violations (ISIv), waveform contamination rate (cR), and presence rate (PR). ISIv is the fraction of spikes that occur within 2ms of the previous spike. cR is the proportion of spikes inside a high-dimensional cluster boundary (by waveform) that are not from the cluster (false positive rate) when setting the cluster boundary at a Mahalanobis distance such that there are equal false positives and false negatives. PR is 1 minus the fraction of 1-minute bins in which there is no spike. We set the following thresholds in qualifying a unit as a single-unit: ISIv < 20%, cR < 0.02, and PR > 90%.

### Analyses

#### Behavioral Analyses

Trials were classified as “attempted” if they did not satisfy the criteria for the other categories: “never started”, “never stopped”, or “aborted”. Never started trials were those where upon trial end (trial maximum duration = 7 seconds), the animal had moved less than 5cm from the origin. Never stopped trials were those where the animals’ linear velocity was above 5cm/s at the of the trial. Aborted trials were those where the animal had started an attempt (linear velocity > 5cm/s) yet travelled less than 30% of the target distance. For the transition probability matrix in **Fig. 1C** and run lengths in **Fig. 2D** rewarded trials were considered “attempted.”

To estimate the reward rate afforded by trials (**Fig. 2E, G,** and **I**), we computed the fraction of rewarded trials for the 30 closest trials within the same session. Similarly, we computed the mean time it took to reach these targets, hence allowing to estimate a reward rate (i.e., probability of reward divided by time). This estimate was robust to the number of adjacent trials used (range tested: 15-60 trials).

To classify sessions as belonging to different animals, we build a linear discriminant analysis (LDA). The model assumes that each class (Y) generates data (X) given a multivariate normal distribution. The covariance matrix of each class is assumed to be the same (empirically estimated, pooled covariance), with the means varying. We use a 10-fold cross-validation, wherein 90% of the data is used to estimate class boundaries, and 10% is used in testing. We iterate over folds to classify all data. The prediction aims to minimize the expected classification cost:

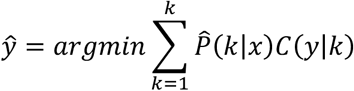

where *ŷ* is the predicted classification, *k* is the number of classes, *P̂*(*k*|*x*) is the posterior probability of class *k* for observation *x* (uniform prior), and *C*(*y*|*k*) is the cost of classifying an observation as y when its true class is k. Th standard cost for LDA was employed.

#### Neural Analyses

Spiking activity was digitized in 1ms bins. Then, within each session (72 in total) each aborted trial was matched to its most similar attempted or aborted trial, by first finding target locations within 5cm (radial) of the target trial. Then, within this subset (typically a dozen trials), we chose the trial with the smallest mean squared error (MSE) in contrasting linear and angular velocity profiles between target and candidate trials. Animals typically followed curvilinear trajectories (see **Fig. 1B**) where the angular component was null or close to null during target presentation. Likewise, given that trials are matched within-sessions, matched by target location and velocity also matches reward rates (i.e., likelihood of attaining a particular reward is computed within sessions).

For single neuron analyses, we performed a spike count within 0 to 300ms from target presentation (i.e., the time-period over which the target was visible), corrected this spike count by trial-specific baseline (-300 to 0 ms), and performed statistical testing comparing matched attempted and aborted trials via a Wilcoxon signed rank test (p <0.05). For visualization, we convolved spike trains with a causal Gaussian kernel (std = 30ms) and then averaged within conditions. To compare fraction of neurons tuned to aborting behavior across brain regions and to chance, we performed Mann–Whitney U tests.

For population-level analysis, we again matched pairs of attempted-aborted or aborted-aborted trials, and expressed the neural activity (averaged between 0 and 300ms post-stimulus onset and corrected for baseline) of *n* neurons in an *n*-dimensional space (**Fig. 3F**). We then computed the Euclidian distance between aborted and attempted trials (*d*_1_), as well as two aborted trials (*d*_2_). We averaged these distances within sessions, and plot histograms of average *d*_1_ and *d*_2_ distances across sessions (i.e., **Fig. 4E**, each count is a session). To quantify the difference in distributions we employ KL divergence. For PCA (**Fig. S6**), we convolved spike trains (as above), then averaged and finally performed PCA. We visualize the first PC, accounting for ∼ 69% of the total variance, and perform statistical testing at each time (from -100 to 1300 ms post-target onset) point via Wilcoxon signed rank test.

### Actor-critic RL models

The actor-critic models used in the current report are augmented version from those presented in Zhang et al., 2024^22^. The formal detail regarding definition of state, action, observation, reward and state transitions are presented here in the *Supplementary Materials.* Similarly, formulas driving the actor and critic updates, and belief modeling, as well as detail to the neural architectures and agent training can be found in the *Supplementary Materials*.

## Acknowledgements

The authors thank Jing Lin and Jian Chen for programming the experimental stimulus. We also thank Eric Avila, Stefania Bruni, and Panos Alefantis for data collection; Roozbeh Kiani for surgical expertise; and Kaushik Lakshminarasimhan for helpful discussions leading to the original RL agents. The work was supported by 1U19NS118246, 1R01NS120407, and 1R01DC014678 from NIH to D.E.A and by K99NS128075 to JPN. JPN was additionally supported by UMN CTSI (1UM1TR004405)/Medical School Early Career Research Award.

## Data and Code Availability

Data are available here: https://osf.io/eybc8/.

Code for RL models is available here: https://github.com/ryzhang1/dlPFC-drives-strategic-aborting

## Supplementary Figures and Figure Captions

**Figure S1.**
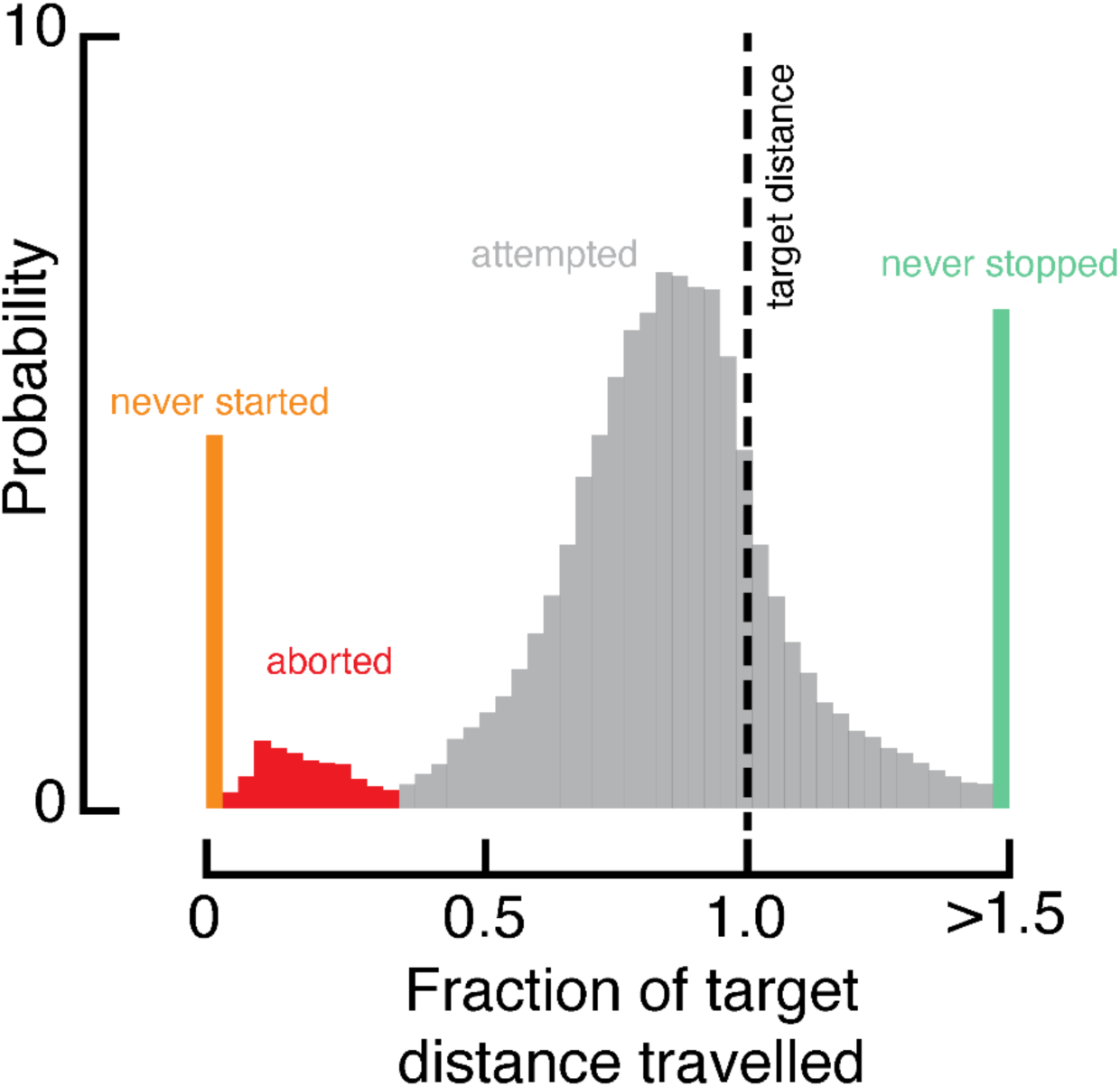
Histogram of macaque stopping distance (radial) expressed as a fraction of target distance. The histogram demonstrates 4 modes; (1) trials wherein the animal did not move at all (“never started”, orange), (2) trials where the animal seemed disengaged with the task and never stopped (green, expressed as travelled distance here but categorized in main text by not stopping for the duration of the trial, 7 seconds), (3) trials where the animal attempted to reach the target (gray), and finally (4) those where the animal stopped well short (<.3 of target distance; red).

**Figure S2.**
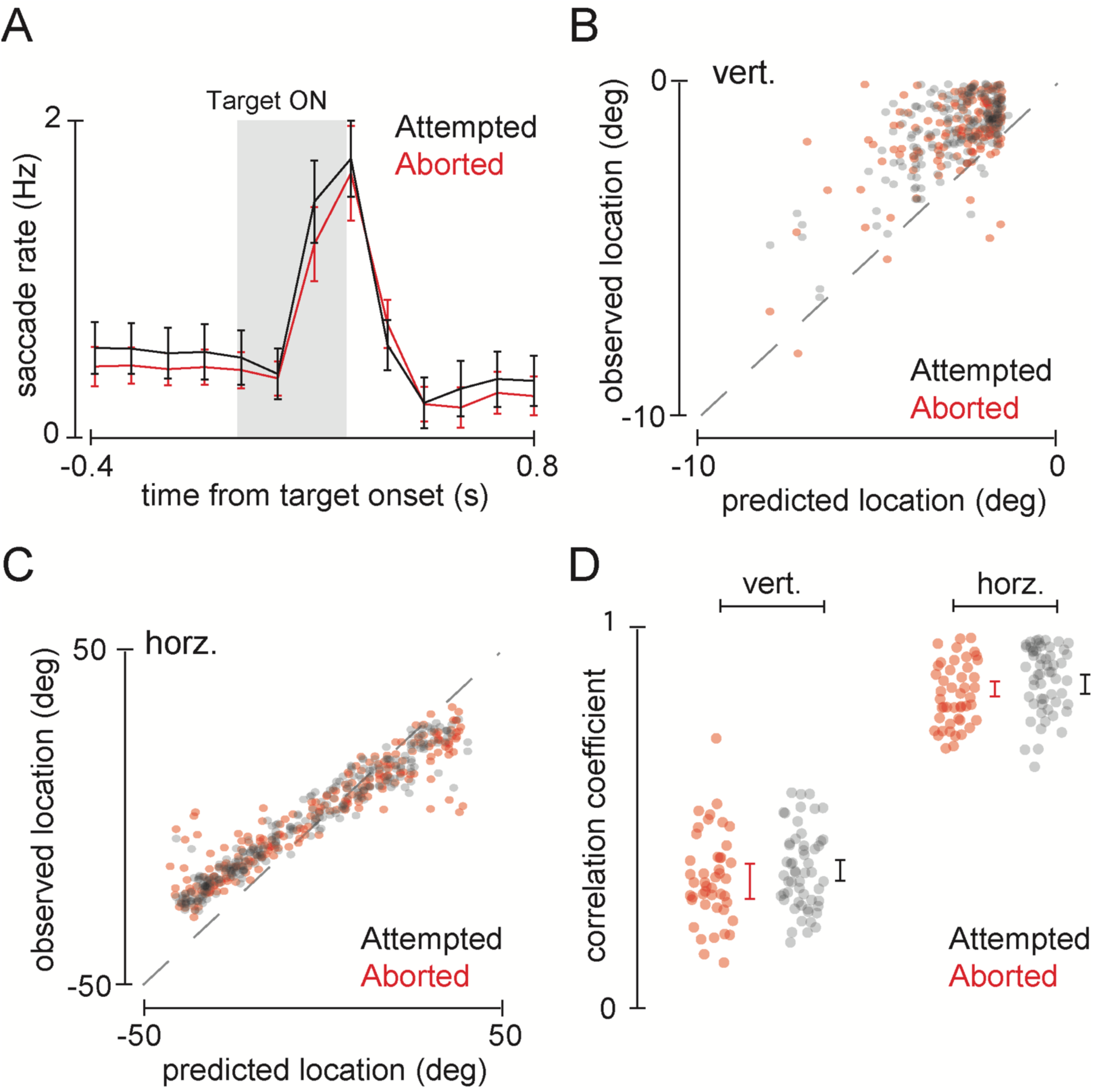
Eye movements during attempted (gray) and aborted (red) trials. **A.** Saccade rate (y-axis) as a function of time from target onset (x-axis) and whether the trial was attempted or aborted. Error bars are ± 1 s.e.m. In both trial types, animals made saccades more frequently shortly after target onset. **B.** Data from an example session showing vertical landing location of saccades following target presentation (y-axis) versus the predicted vertical location if animals were saccading toward the target. During both attempted (gray) and aborted (red) trials, the animals appear to saccade toward the target. **C.** Data from an example session (same as **B.**) showing horizontal landing location of saccades following target presentation (y-axis) versus the predicted vertical location if animals were saccading toward the target. During both attempted (gray) and aborted (red) trials, the animals appear to saccade toward the target. **D.** Summary across all sessions of the correlation coefficient of data plotted in **B** (left panel) and **C (**right panel). Each dot is a session, error bars show ± 1 s.e.m around the mean. Overall, saccades are more accurate along the horizontal than vertical dimension. Importantly, the correlation between predicted and observed saccade landing was not different across attempted (gray) and aborted (red; all p > 0.78) trials.

**Figure S3.**
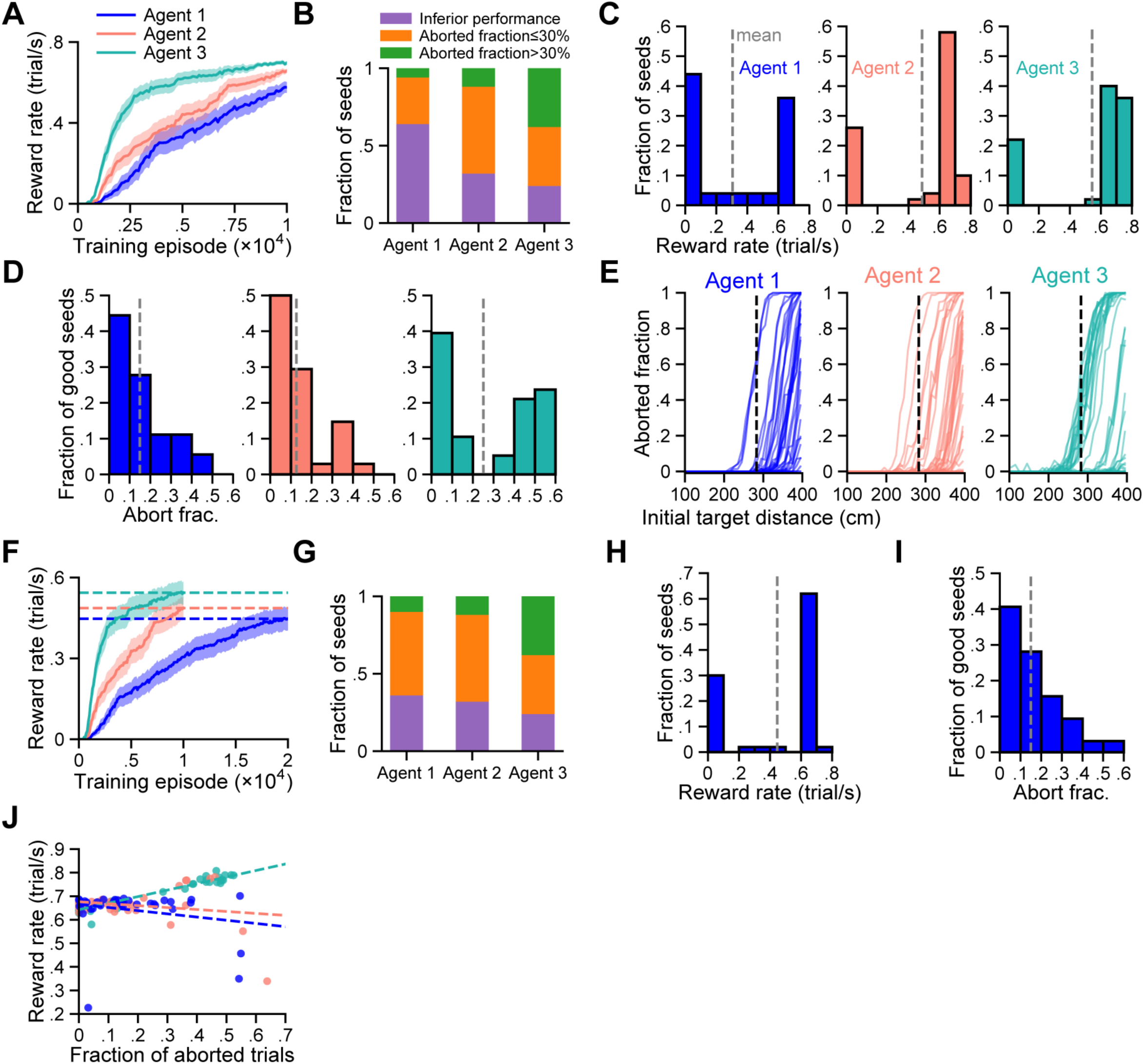
Supplementary figures for Fig. 2. **A.** Similar to Fig. 2F, but excluding non-learning seeds (reward rate < 0.2). **B.** Fraction of seeds with a reward rate ≤ 0.6 (inferior performance), and among the remaining seeds, the fraction that aborted ≤ 30% or > 30% of trials. **C.** Distribution of the reward rate for all random seeds for each agent. 500 trials were used to evaluate each seed. Gray dashed lines denote the means. **D.** Similar to (C), but showing the fraction of aborted trials for random seeds with a reward rate > 0.6. **E.** Similar to Fig. 2H, but showing data for each random seed. **F.** Similar to Fig. 2F, but Actor 1 (blue) was trained twice as long. Dashed lines denote the performance after training. **G, H, I, and J.** Similar to B, C (left), D (left), and Fig. 2I, but Agent 1 was trained for twice as long as before.

**Figure S4.**
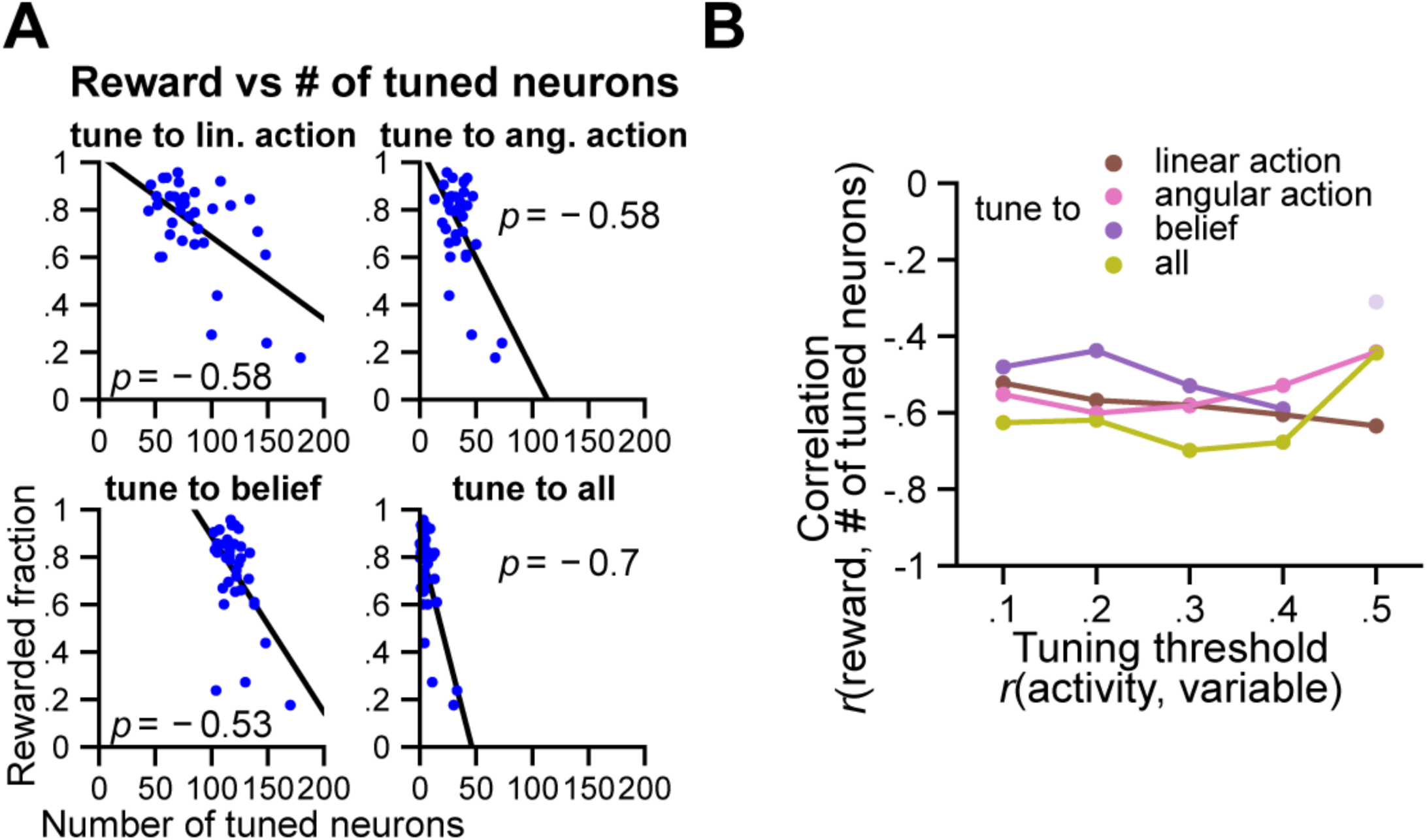
Good performance is associated with greater neural specialization in Actor 1. **A.** Fraction of rewarded trials versus number of neurons tuned to linear action, angular action, target distance (belief), or all three variables for Agent 1. For each neuron, an absolute correlation between its activity and the investigated variable(s) higher than a tuning threshold of 0.3 is considered tuned. Each dot denotes a random seed. 2000 trials were tested for each seed. Agent 1 was trained as in Fig. S3G. **B.** Showing the correlation between rewarded fraction and number of tuned neurons across random seeds in A (tuning threshold = 0.3), and also including results using additional thresholds. Line-connected dots indicate significance (*p* < 0.05), while the dim, unconnected one do not.

**Figure S5.**
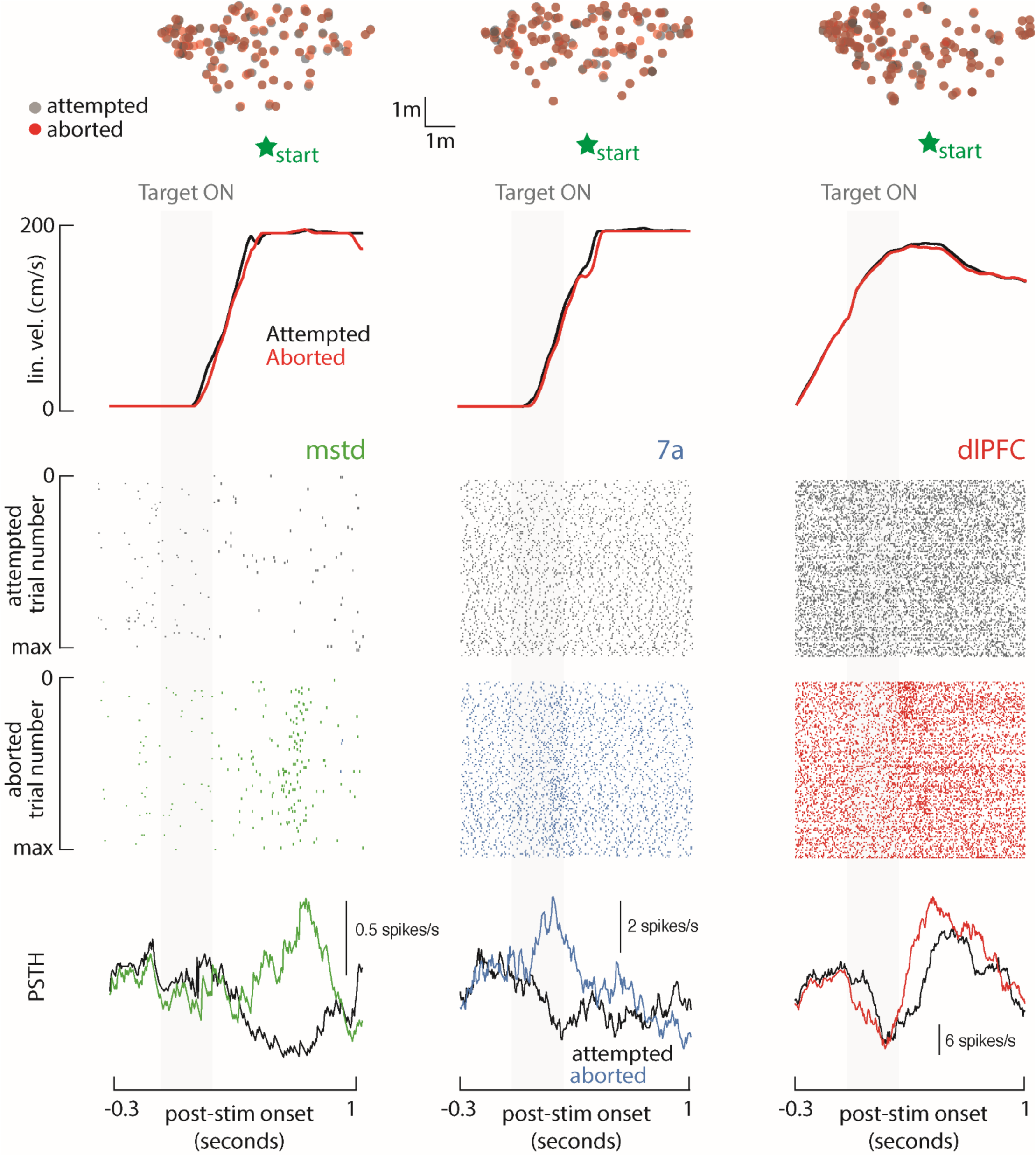
Additional examples of single units differentiating between matched aborted (black) and attempted trials (colored). The figure follows the format from Fig. 4A, demonstrating the heterogeneity in responses observed.

**Figure S6.**
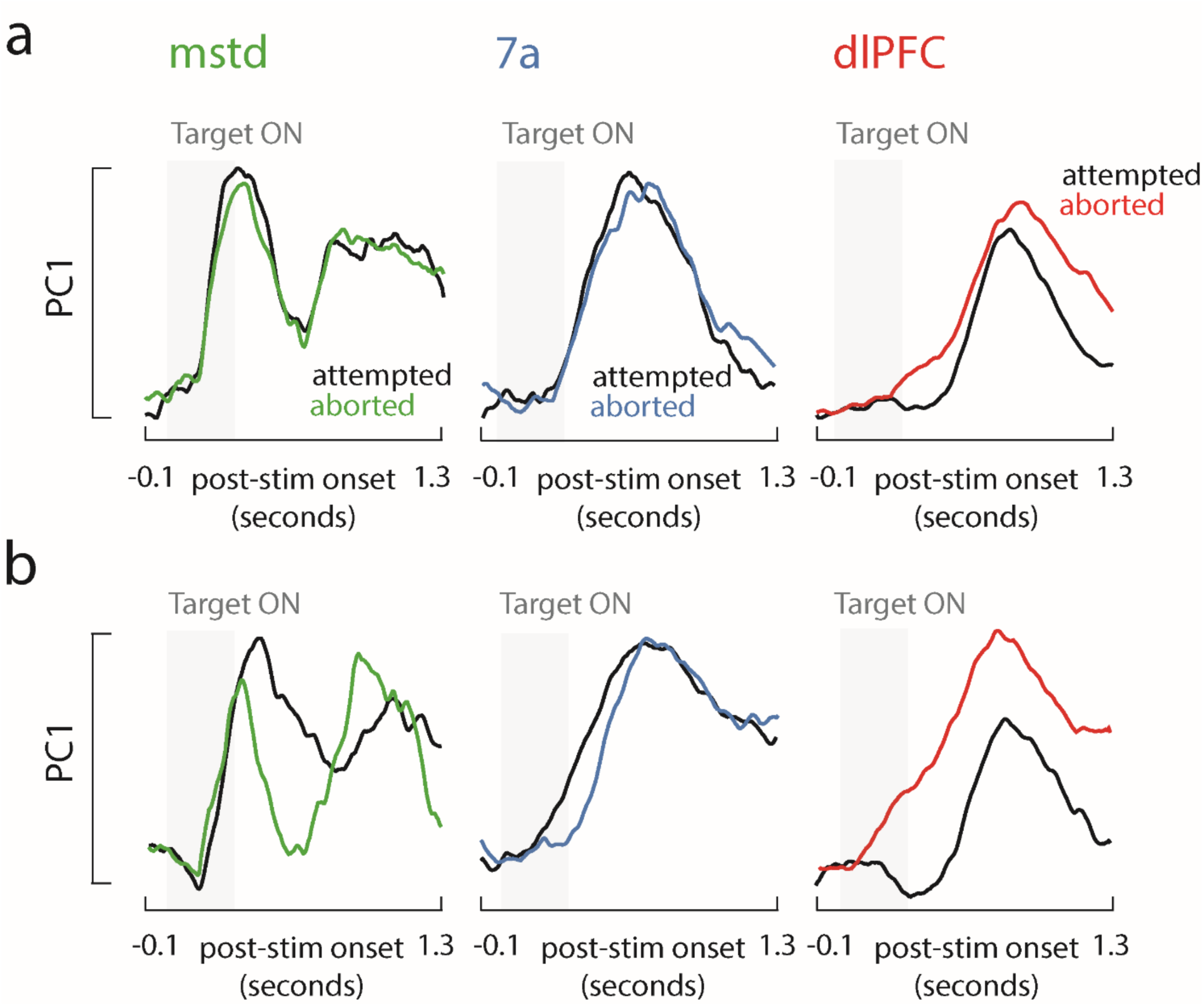
Principle component analysis examining population dynamics preceding aborted (colored) or attempted (black) trials. **A.** First, we include all neurons in this analysis – both those significantly and not differentiating between attempted and aborted trials. We plot the principal component accounting for most variance in each area (68.4% of the total variance on average) as a function of time from stimulus onset. The results show robust evoked responses in all areas, regardless of whether the animals subsequently aborted the trial or not. Most importantly, mimicking the single cell results (Fig. 4A**-D**), this analysis showed a divergence in the evoked response to subsequently attempted and aborted trials in dlPFC (p < 0.05 between 291ms and 639ms post-stimulus onset, red), but not MSTd (green) or 7a (blue, all p > 0.05). **B.** When including in this analysis only cells distinguishing between attempted and aborted trials, the evoked responses diverged first in dlPFC (114ms post-stimulus onset, red), then in 7a (185ms, blue), and finally in MSTd (300ms, green).

**Figure S7.**
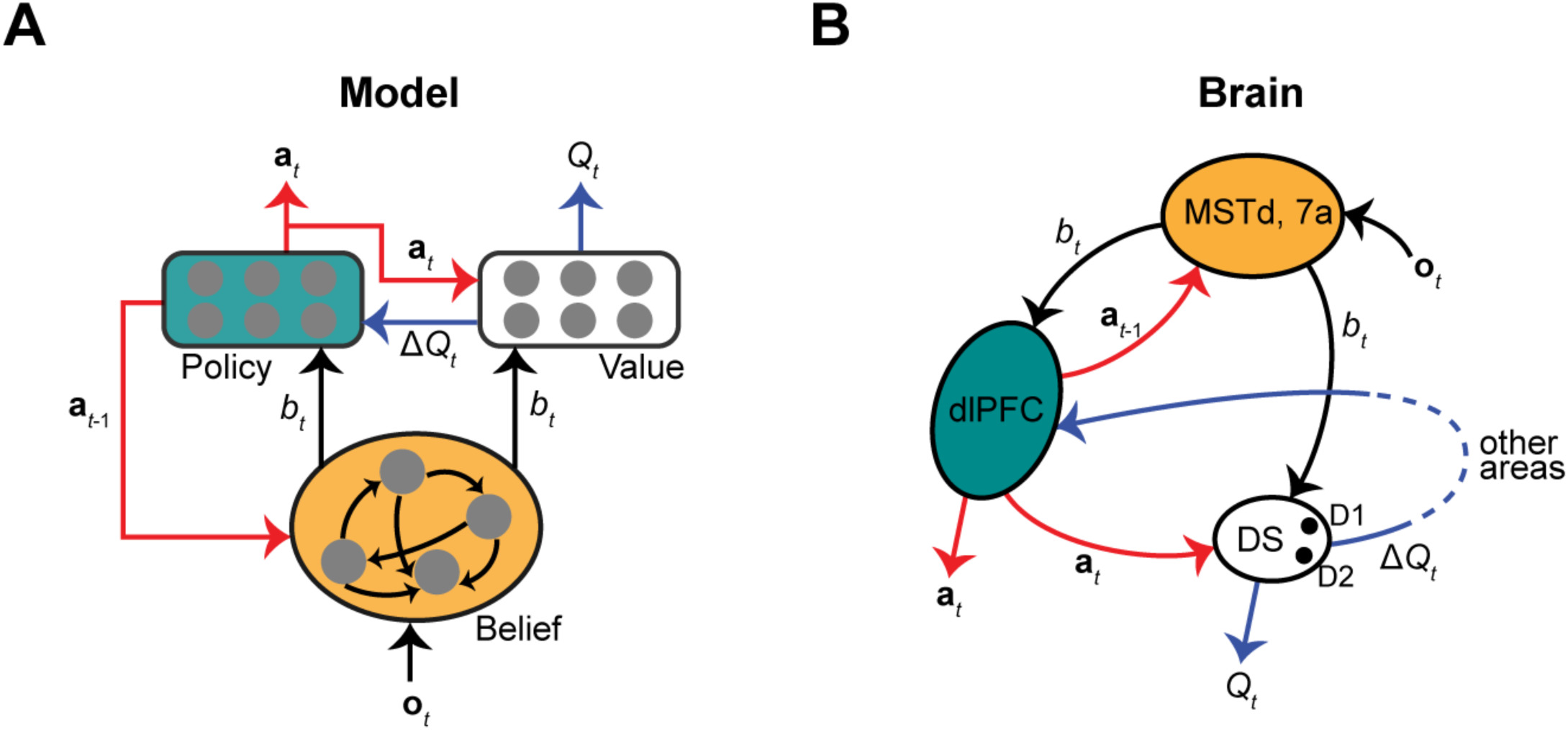
Analogy between our model and the prefrontal-striatal system. **A.** Schematic of our model, similar to Fig. 3D, but showing only input and output arrows. Red and blue arrows denote projections from the policy and value modules, respectively. **B.** A hypothetical prefrontal-striatal circuit implementing our model. DS: dorsal striatum. D1 and D2: two types of dopamine receptors. Dashed blue arrow denotes indirect connections, as the DS projects to the dlPFC indirectly through other areas.

